# Computational proteogenomic identification and functional interpretation of translated fusions and micro structural variations in cancer

**DOI:** 10.1101/168377

**Authors:** Yen Yi Lin, Alexander Gawronski, Faraz Hach, Sujun Li, Ibrahim Numanagić, Iman Sarrafi, Swati Mishra, Andrew McPherson, Colin Collins, Milan Radovich, Haixu Tang, S. Cenk Sahinalp

**Affiliations:** School of Computing Science, Simon Fraser University, Burnaby, BC, Canada; School of Informatics and Computing, Indiana University, Bloomington, IN, USA; Vancouver Prostate Centre, Vancouver, BC, Canada; Dept. of Urologic Sciences, University of British Columbia, Vancouver, BC, Canada; Department of Surgery, Indiana University, School of Medicine, Indianapolis, IN, USA

## Abstract

**Motivation:** Rapid advancement in high throughput genome and transcriptome sequencing (HTS) and mass spectrometry (MS) technologies has enabled the acquisition of the genomic, transcriptomic and proteomic data from the same tissue sample. In this paper we introduce a novel computational framework which can integratively analyze all three types of omics data to obtain a complete molecular profile of a tissue sample, in normal and disease conditions. Our framework includes MiStrVar, an algorithmic method we developed to identify micro structural variants (microSVs) on genomic HTS data. Coupled with deFuse, a popular gene fusion detection method we developed earlier, MiStrVar can provide an accurate profile of structurally aberrant transcripts in cancer samples. Given the breakpoints obtained by MiStrVar and deFuse, our framework can then identify all relevant peptides that span the breakpoint junctions and match them with unique proteomic signatures in the respective proteomics data sets. Our framework's ability to observe structural aberrations at three levels of omics data provides means of validating their presence.

**Results:** We have applied our framework to all The Cancer Genome Atlas (TCGA) breast cancer Whole Genome Sequencing (WGS) and/or RNA-Seq data sets, spanning all four major subtypes, for which proteomics data from Clinical Proteomic Tumor Analysis Consortium (CPTAC) have been released. A recent study on this dataset focusing on SNVs has reported many that lead to novel peptides [1]. Complementing and significantly broadening this study, we detected 244 novel peptides from 432 candidate genomic or transcriptomic sequence aberrations. Many of the fusions and microSVs we discovered have not been reported in the literature. Interestingly, the vast majority of these translated aberrations (in particular, fusions) were private, demonstrating the extensive inter-genomic heterogeneity present in breast cancer. Many of these aberrations also have matching out-of-frame downstream peptides, potentially indicating novel protein sequence and structure. Moreover, the most significantly enriched genes involved in translated fusions are cancer-related. Furthermore a number of the somatic, translated microSVs are observed in tumor suppressor genes.

## 1 Introduction

Rapid advances in high throughput sequencing (HTS) and mass spectrometry (MS) technologies has enabled the acquisition of the genomic, transcriptomic and proteomic data from the same tissue sample. The availability of three types of fundamental omics data provide complementary views on the global molecular profile of a tissue under normal and disease conditions [2]. Recently developed computational methods have aimed to integrate two or three of these data types to address important biological questions, such as (i) correlating the abundances of transcription and translation products [3]; (ii) detecting peptides associated with un-annotated genes or splice variants (in mouse [4], C. elegans [5], zebrafish [6] and human samples [7, 8]); (iii) characterizing chimeric proteins by searching unidentified tandem mass spectrometry(MS/MS) data through the use of conventional peptide identification algorithms applied to a pre-assembled database of “known” chimeric transcripts from the literature [9].

In the past year or so, several studies have aimed to identify novel peptides matching patient specific transcripts derived from RNA-Seq data. For example, Zhang et al. [10] focused on identifying novel peptides involving Single Amino Acid Variants (SAAVs) in colorectal cancer. A later study by Cesnik et al. [11] also considered novel splice junctions and (a limited set of user defined) Post-Translational Modifications (PTM) in a number of cell lines. Because of the importance of phosphorylation in cellular activity and cancer treatment [12], this was further expanded to identify novel phosphorylation sites by Mertins et al. [1], on the CPTAC breast cancer data set, which is the subject of our paper. However, none of these studies aimed to perform integrative analysis of transcribed and translated genomic structural alterations such as fusions, inversions and duplications in tumor tissues.

**Genomic structural variants (SVs)** alter the sequence composition of associated genomic regions in a significant manner. Major SV types include (segmental) deletions, duplications (tandem or interspersed), inversions, translocations and transpositions. SVs observed in exonic regions may lead to aberrant protein products. Many such SVs have been associated with disease conditions and especially cancer. Common SVs associated with cancer include deletions in tumor suppressors such as BRCA1/2 [13] in breast cancer, duplications in FMS-like tyrosine kinase (FLT3) gene in acute myeloid leukemia (AML) [14] and an inversion causing cyclin D1 overexpression in parathyroid neoplasms [15].

A **gene fusion** occurs when exonic regions of two (or more) distinct genes are concatenated to form a new chimeric gene, as a result of a large scale SV. Gene fusions can disrupt the normal function of one or both partners, for example by up-regulating an oncogene (e.g. TMPRSS2-ERG) or generating a novel or truncated protein (e.g. BCR-ABL1 [16]). They have been demonstrated to play important roles in the development of haematological disorders, childhood sarcomas and in a variety of solid tumors. For example, ETS gene fusions are present in 80% of malignancies of the male genital organs, and as a result these fusions alone are associated with 16% of all cancer morbidity [17]. Others, including the EML4-ALK fusion in non-small-cell lung cancer and the ETV6-NTRK3 fusion in human secretory breast carcinoma occur in much lower frequency [18, 19]. The discovery of such low-recurrence gene fusions may be of significant clinical benefit since they have potential to be used as diagnostic biomarkers or as therapeutic targets - if they encode novel proteins affecting cancer pathways [20–22].

There are a number of available computational tools for detecting structural variants, each based one or more of the following general strategies. (1) Detection of variants using discordantly mapping paired end reads, more specifically read mappings that either invert one or both of the read ends, or change the expected distance between the read ends. Tools using this approach include Breakdancer [23] and VariationHunter [24]. (2) Detection of variants using split-read mappings - which partition a single end read into two and map them independently to two distant loci - or soft-clipped read mappings - which map only a prefix or suffix of a read. One example employing this approach is Socrates [25]. (3) Detection of variants using an assembly based approach. These tools map assembled contigs for improved precision. Examples include Barnacle [26] and Dissect [27] (both of which happen to be RNA-Seq analysis tools, but can also be used to analyze genomic data). Additional tools employing a combination of these strategies include Pindel [28], Delly [29], GASVPro [30] and HYDRA [31].

Our focus in this paper is microSVs (micro structural variants), i.e. events involving genomic sequences shorter than a few hundred bps, especially in exonic regions, since they are more likely to result in a translated protein. Available tools for SV discovery typically fail to capture microSVs, or do so while producing many false positives, thus the problem of robustly discovering microSVs remain open.

In contrast to microSVs, gene fusions can be inferred at a large scale by detecting chimeric transcripts in RNA-Seq data [32]. Currently, there are two general computational approaches to detect gene fusions. (i) The mapping-based approach (e.g. deFuse [33], FusionMap [34], FusionSeq [35], ShortFuse [36], SOAPfuse [37], and TopHat-Fusion [38]) suggests to first map RNA-Seq reads to the reference genome, and then discover fusion transcript candidates by analyzing discordant mappings. More involved methods in this category include nFuse [39] and Comrad [40], which incorporate WGS (Whole Genome Sequencing) data for more accurate predictions and handling complex fusion patterns that involve three or more genes. (ii) The assembly-based approach such as Barnacle [26] and Dissect [27], on the other hand, suggests to first *de novo* assemble RNA-seq reads into longer contigs by using available transcriptome assemblers (e.g., Trinity [41]), and only then map the assembled contigs back to the reference genome, with the aim of reducing the potential errors introduced by mapping short reads to the reference genome.

**Our first contribution** in this paper is a novel algorithmic tool named **MiStrVar** (**Mi**cro **Str**uctural **Var**iant caller), which identifies microSV breakpoints at single-nucleotide resolution by (1) identifying each one-end-anchor (OEA), i.e. a paired-end read where one end maps to the reference genome and the other end cannot be mapped, (2) clustering OEAs based on (i) mapping loci similarity and (ii) the possibility of assembling the unmappable ends into a single contig, and (3) aligning the contig formed by unmappable ends with the reference genome - in the vicinity of the mapped ends - simultaneously detecting putative inversions, duplications, indels or single nucleotide variants (SNVs) through a unified dynamic programming formulation.

MiStrVar approach has several advantages over existing SV discovery tools. Firstly, MiStrVar analyzes many more reads than those considered by the tools using only split-reads or soft clipped reads. Any mapped read which has a hamming distance to the reference greater than four (as a default parameter, which can be user modified) is considered for assembly. This allows for the discovery of inversions or duplications as short as 5bp and inversions with palindromic sequences, improving sensitivity. Secondly, this approach is much less time consuming than assembly based methods, since only the subset of unmappable reads are assembled rather than the entire genome. Finally, MiStrVar uses a unified dynamic programming formulation, superior to tools that identify each type of variant individually, especially because these tools misinterpret certain variants, such as inversions, as a combination of other variants. See Supplementary Figure 1 for a detailed illustration.

Both fusions and microSVs may be independently observed in genomic, transcriptomic, and proteomic data; however, the most impactful aberrations, especially in the context of cancer, are the ones that can be observed in all levels in the same tissue simultaneously. In such cases, integrative analysis of these three omics data types can provide independent evidence for the presence and heritability of aberrations. For example, transsplicing events, which lead to chimeric transcripts, can only be observed in transcriptomic (but not in genomic) data, and thus can be distinguished from fusion events with genomic breakpoints through simultaneous analysis of genomic and transcriptomic data acquired from the same sample.

The vast majority of large-scale studies of sequence aberrations are based on genomic and transcriptomic data. Most proteogenomics research mainly focuses on detecting single amino acid variants and studying protein abundances affected by single nucleotide variants [10, 42]. No available large-scale study has been conducted on the detection and validation of aberrant proteins and their genomic and transcriptomic origins. As mentioned earlier, expressed aberrant genome variants can have considerable functional influence on proteins, and as such, they may affect molecular pathway activity or pathogenesis in disease, especially in cancer. Detection of aberrant protein variants provides new insights into diagnostic marker identification and drug development (recurrent protein aberrations can imply potential drug targets) and can help develop novel strategies for therapeutic intervention.

Proteomic technologies have enabled high throughput, sensitive and deep protein analysis for complex disease-associated samples, aiming at discovering potential disease protein biomarkers [43–45], including low-abundant proteins or protein isoforms, or variants. Moreover, proteomic analyses can provide complementary information to transcriptomic and genomic analysis, as proteomic analyses are carried out by completely different technologies (i.e., mass spectrometry or MS) from DNA sequencing. Furthermore, advancement in MS instrumentation has enabled proteomic analysis to achieve sensitivity on par with RNA-seq in detecting low abundant events of gene expression in complex samples [10]. Therefore, integrating transcriptomic and proteomic data can improve both the sensitivity and confidence in characterizing expressed aberrant variants in complex samples such as tumor tissues.

**Our second contribution** in this paper is **ProTIE** (**ProT**eogenomics **I**ntegration **E**ngine), the first computational framework that integrates high throughput genomic, transcriptomic and proteomic data to identify translated structural aberrations, specifically gene fusions and microSVs, in protein-coding genes. In particular, ProTIE takes sequence aberrations from WGS and RNA-Seq data as its input and validates them on the mass-spectrometry based proteomics data, while ensuring that each such proteomic signature is unique to the matching sequence aberration. By integrating multiple data sources simultaneously, ProTIE is able to provide a strongly supported set of candidate aberrations from the highly sensitive results of MiStrVar and deFuse. This is particularly helpful for selecting target events or genes for clinical studies.

**Results.** We ran our computational framework to detect all translated gene fusions in RNA-Seq (low coverage 50bp paired-end) data in the complete set of 105 TCGA (The Cancer Genome Atlas) breast cancer samples for which CPTAC (Clinical Proteomic Tumor Analysis Consortium) mass spectrometry data have been released.^1^ These 105 samples include all four of the most common intrinsic subtypes of breast cancer. Among them, 22 samples also have matching WGS data, on which we used our framework to identify exonic microSVs. This resulted in 206,255 fusions and 69,876 microSVs across the 105 samples. 2,215 of these microSVs are also supported by transcriptomic (RNA-Seq) evidence.

All breakpoints from the predicted fusions and microSVs were then analyzed for identifying supporting peptides from mass spectrometry data. This yielded 244 aberrant peptides from 432 possible aberrations. More specifically, 169 novel peptides originate from 295 fusion candidates (many of the fusions are recurrent and thus produce the same novel fusion peptide) and 75 peptides originate from 137 potential microSVs; this is of particular note since many of the genomic microSVs are recurrent, yet the ones that are translated are mostly private. Note that a sequence aberration may give rise to more than one novel peptide in case it results in a frameshift. See Table 1 for a summary of results. ^2^

**Table 1:**
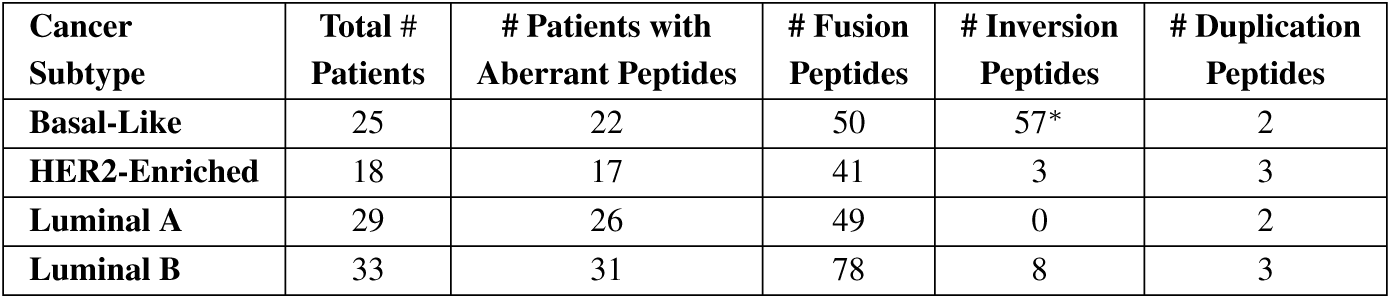
Distribution of 244 detected, high confidence, aberrant peptides over four breast cancer subtypes, across 105 patients. **# Patients with aberrant peptides** indicate the number of patients with either detected fusion peptides or microSV peptides in that subtype. As can be seen, all but one of the patients exhibit at least one translated fusion or microSV. The next three columns respectively indicate the number of peptides detected from fusions, microinversions and microduplications, within specific subtypes. *The high number of microinversion peptides in Basal-Like breast cancer can be attributed to two patients, A0CM, A0J6, whose genomes had gone through substantial reorganization.

## 2 Methods

Our computational framework (see Figure 1), is comprised of a number of algorithmic tools that we developed for detecting transcriptomic and genomic aberrations, and searching for expressed protein variants resulting from these aberrant sequences. Given a set of genomic (WGS), transcriptomic (RNA-seq) and proteomic (Mass Spectrometry) data, each collected from the tumor tissue of a patient, our pipeline detects translated *sequence aberrations* in three major steps.

1. Each whole genome sequencing dataset is analyzed with MiStrVar, the microSV discovery tool we introduce in this paper, to identify microSVs occurring in proteincoding genes. (Note that our computational framework provides the option of validating genomic microSVs at the transcriptomic level by identifying RNA-Seq reads associated with each microSV breakpoint.)
2. Each transcriptomic dataset is analyzed by our in-house fusion detection method deFuse [33], which reports potential fusion events between two protein coding genes, and the *fused* transcript sequences spanning the fusion breakpoints. (Note that our computational framework enables the use of our integrative fusion detection methods nFuse [39]/Comrad [40] for corroborating potential fusions observable in WGS and RNA-Seq data.)
3. All omics data is finally integratively analyzed through ProTIE, our novel ProTeogenomics Integration Engine as follows. Each mass spectrometry dataset is searched against a protein sequence database consisting of all human proteins from Ensembl human protein database GRCh37.70 [46], along with a database of proteins generated by fused transcripts and microSVs, by the use of MS-GF+ search engine [47]. Aberrant peptides identified by the procedure with high confidence (e.g., at 1% false discovery rate estimated by using the target-decoy approach [48]) are reported, provided they are also detected in the genomic/transcriptomic dataset from the same tumor tissue sample. (For further validating aberrations identified at multiple omics levels, our computational framework also provides the option of searching for recurrences across multiple tumor samples, possibly representing the same tumor subtype.)

**Figure 1:**
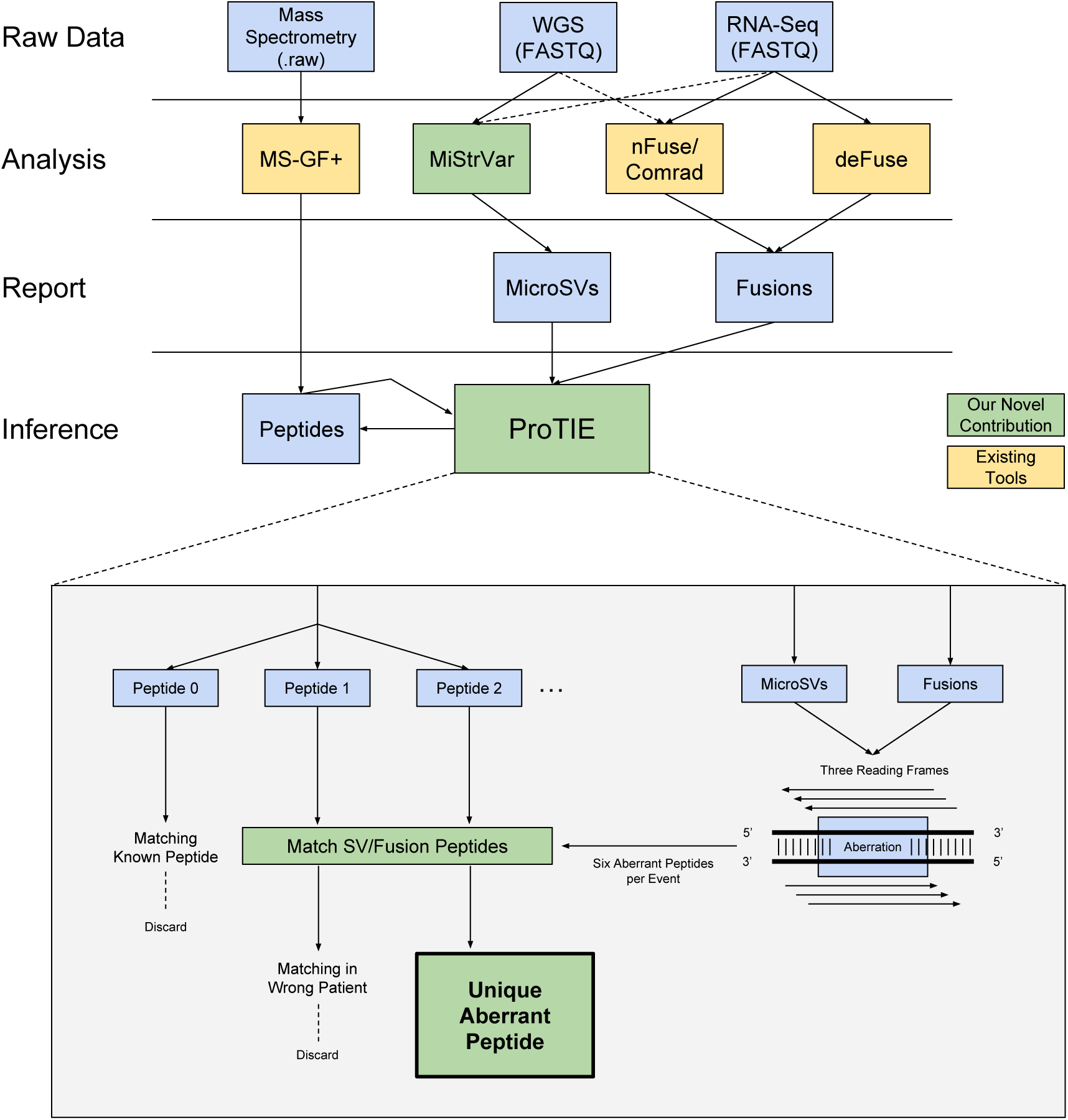
Overview of the computational pipeline for identifying translated sequence aberrations. Mass spectrometry data is used to validate fusions detected in the RNA-Seq data and micros Vs detected in the WGS data. For tumor samples with matching RNA-Seq and WGS data, our pipeline provides the ability to detect transcribed micros Vs and fusions with genomic origins as well. The pipeline introduces MiStrVar, a tool for detecting micros Vs from WGS data. It also features our in house developed fusion discovery tool(s) deFuse (as well as nFuse/Comrad), as well as the MS-GF+ mass spectroscopy search engine. The final step is the ProTIE (Proteogenomics integration engine) for sequence and mass spectrometry data: After running deFuse and MiStrVar to respectively identify fusions and microSVs, we generate each possible breakpoint peptide from the 6 distinct reading frames associated with each of these aberrations. For mass spectra from the same tumor sample, we discard those which can be matched to known proteins, and keep only spectra matched to breakpoint peptides identified above. The resulting high quality peptide-spectra matches (PSM) provide proteomics-level evidence for the predicted aberrations.

### 2.1 Detection of Fusions and microSVs in WGS and RNA-Seq Data

To detect fusions in RNA-Seq data, we applied deFuse [33] which predicts fusion transcripts based on analyzing discordantly mapped read-pairs and one-end anchors. To detect microSVs in WGS data, we applied our novel micro-structural variant caller, MiStrVar, which works in three major steps (See Figure 2 in Supplementary materials for an overview):

In **step (A)**, MiStrVar identifies all one-end anchors (OEA) in the read data: an OEA is a paired-end-read for which only one end maps to the reference genome within a user defined error threshold. Once all reads are (multiply) mapped to a reference genome using mrsFAST-ultra [49, 50], and all OEAs are extracted, the mapped ends of OEAs are clustered based on the mapping loci. MiStrVar provides the user two options for cluster identification, each satisfying one of the following distinct goals. For applications where sensitivity is of high priority, MiStrVar employs a sweeping algorithm for OEA mapping loci (introduced for VariationHunter [24]). For applications where running time is of high priority, MiStrVar employs an iterative greedy strategy.

In **step (B)**, for each OEA cluster identified in step (A), MiStrVar assembles the unmapped end of the reads to form contigs (of length <400bp in practice) by aiming to solve the NP-hard [51] **dominant superstring (DSS)** problem. MiStrVar employs a greedy strategy similar to that used to compute a constant factor approximation to the shortest superstring problem [52].

In **step (C)**, each contig associated to an OEA cluster is aligned to a region (of length several kilobases long) surrounding the OEA mapping loci, first through a simple *local*-*to*-*global* sequence alignment algorithm, that does not consider any structural alteration. (The reverse complement of the contig is also aligned to the same region.) The start and end position of this first, crude alignment is used to determine the approximate locus and length of the potential microSV implied by the contig. The exact microSV breakpoints are obtained in the next step through a more sophisticated alignment that considers structural alterations, which is applied to the portion of the reference genome restricted by the first alignment. The dynamic programming formulation for this alignment is an extension of the Schöniger-Waterman algorithm [53] which was designed to capture inversions in the alignment. Specifically, the extensions enable the user to

1. discover the single best optimal event, rather than an arbitrary number of events,
2. handle gaps extending over breakpoints (in cases of missing contig sequence), and,
3. simultaneously predict duplications, insertions, deletions and SNVs in addition to inversions.

Further details on the methodology of deFuse and MiStrVar can be found in the supplementary text.

### 2.2 Identification of Translated and Transcribed Sequence Aberrations

ProTIE provides the ability to detect translated aberrations by searching mass spectra against an aberrant peptide database. More specifically, given transcriptomic breakpoints pointing to fusions or microSVs, ProTIE identifies respective aberrant peptides from proteomic data by first generating a peptide database, and then identifying aberrant peptides based on mass spectrometry search results provided by MS-GF+ [47]. (See subsection 2.3 in supplementary materials for details of database construction and parameters used in proteomics search.)

Our pipeline also provides the user with the additional ability to jointly analyze matching WGS and RNA-Seq data for identifying transcribed genomic (in fact genetic) microSVs. Given a set of genomic microSVs, along with their breakpoints detected by MiStrVar, our pipeline generates corresponding aberrant transcripts. It then maps RNA-Seq reads to the collection of these aberrant transcripts. After filtering reads that can be mapped to a known isoform or potential novel spliceform, the remaining mappings provide evidence for aberrations in transcribed regions. See subsection 2.4 in supplementary materials for details about mappings and read filtration steps.

### 2.3 Availability

MiStrVar is available for download at https://bitbucket.org/compbio/mistrvar, and ProTIE is available at https://bitbucket.org/compbio/protie.

## 3 Experimental Results

*CPTAC Breast Cancer Dataset*. Clinical Proteomic Tumor Analysis Consortium (CPTAC, http://proteomics.cancer.gov) [54, 55] aims to provide proteogenomic characterization of specific cancers based on joint analysis of proteomic, transcriptomic, and genomic data acquired from the same group of cancer patients. CPTAC currently focuses on the relationship between protein abundance, somatic mutations and copy number alterations occurring in cancer-related genes [10]. Information about aberrations hidden in unidentified spectra and unmapped sequenced reads have not been revealed in the current CPTAC analysis framework; this happens to be the main focus of our paper.

At the time of submission of this paper, proteomics data for tumor samples from three cancer types had been released by CPTAC: colorectal cancer, breast cancer, and ovarian cancer. In addition, The Cancer Genome Atlas (TCGA, http://cancergenome.nih.gov/) has released RNA-Seq and WGS data on both normal and tumor tissues from the same group of patients through Cancer Genomics Hub (CGHub, https://cghub.ucsc.edu/). RNA-Seq data for breast and ovarian cancer patients are in the form of paired-end reads, however, for most of colon and rectal cancer samples only singleend reads were collected. Because we rely on paired-end mappings for detecting fusions and microSVs and since the RNA-Seq data from normal tissues from the ovarian cancer patients had not been released at the time of the submission, our focus in this paper is the breast cancer dataset. Details about CPTAC samples used in our analysis can be found in Supplementary Tables 4, 5.

*Breast Cancer Cell Line*. In addition to the CPTAC and TCGA datasets, we used the HCC1143 ductal breast cancer cell line (triple negative breast cancer cell line from ATCC) for which we obtained matching tumor/normal Illumina HiSeq WGS, RNA-Seq and mass spectrometry data. The matching normal cell line, HCC1143-BL, is a B lymphoblastoid cell line initiated from peripheral blood lymphocytes from the same patient as HCC1143 by transformation with Epstein-Barr virus (EBV). The WGS data was obtained from NCI Genomic Data Commons (https://gdc.cancer.gov/), originally sequenced as part of the Cancer Cell Line Encyclopedia Project [56]. We used this cell line as preliminary validation for our approach before starting full scale analysis.

### 3.1 Gene Fusion Detection by deFuse

*Gene Fusions in the HCC1143 Breast Cancer Cell Line*. We have run our fusion detection method, deFuse to detect gene fusions on RNA-Seq data from HCC1143 cell line. There are 81.73M paired end reads of 101bp length. Based on concordant mapping results, the average fragment length and standard deviation were 264.2bp and 86.59 bp respectively. deFuse predicts 1,325 fusions from this dataset, out of which 74 are considered high confidence predictions based on the filtering criteria employed by deFuse [33].

*Gene Fusions in Breast Cancer Patient RNA-Seq Data*. Each RNA-Seq dataset from the CPTAC breast cancer patient cohort was, on average, comprised of 76M paired-end Illumina reads with length 50bp. Based on transcriptome mapping results, the average fragment length and standard deviation were 190.3bp and 65.47bp respectively. In total deFuse detected 206,255 fusions; on average, this amounts to 1,964 predictions per sample. However, many of these predictions had low deFuse scores, either due to low sequence similarity or limited read support, and thus were not good fusion candidates. Only 3,907 of these predictions (roughly 2% of all predictions) in total are considered to be high confidence calls by deFuse.

### 3.2 MicroSV Detection by MiStrVar

MicroSV predictions were based on three WGS datasets. The first is a simulation dataset based on the Venter genome developed with the goal of assessing sensitivity and precision of our methods with respect to available tools for SV discovery. These results are summarized in Table 2; more details can be found in supplementary materials. The second dataset consists of WGS data from the HCC1143 cell line (both tumor and normal), which was used to assess our methods’ accuracy on a homogeneous tumor sample. The third dataset is comprised of 22 TCGA/CPTAC breast cancer WGS data, which were used for full scale evaluation of our methods.

**Table 2:** Comparison of precision, recall, false discovery rate (FDR) and false negative rate (FNR) of MiStrVar against other SV discovery tools. All tools were run with default parameters and the calls for each microSV type (we only considered the calls made by each tool for that microSV) were called true or false based on the metrics provided by the tools (quality, identity or support, if they exist). The threshold values for each metric were chosen to maximize the F-score (Supplementary Table 1). Only inversions of length ≤400bp were considered in the calculations. If a tool does not provide precise breakpoints, breakpoints falling within a provided range are counted as true positives. Known insertion SNPs were filtered for all duplication results.

| SV Type | Tool | 5-100 bp |  |  |  | 101-400 bp |  |  |  |
| --- | --- | --- | --- | --- | --- | --- | --- | --- | --- |
|  |  | Precision | Recall | FDR | FNR | Precision | Recall | FDR | FNR |
| Inversions | MiStrVar | 91.20% | 92.68% | 8.80% | 7.32% | 93.10% | 98.78% | 6.90% | 1.22% |
|  | Breakdancer | 66.67% | 1.63% | 33.33% | 98.37% | 59.00% | 95.00% | 41.35% | 4.88% |
|  | Delly | 67.00% | 1.63% | 33.00% | 98.37% | 61.98% | 91.46% | 38.02% | 8.54% |
|  | Pindel | 82.64% | 81.30% | 17.36% | 18.70% | 88.51% | 93.90% | 11.49% | 6.10% |
|  | SoftSV | 0.00% | 0.00% | 100.00% | 100.00% | 93.75% | 18.29% | 6.25% | 81.71% |
| All Duplications | MiStrVar | 30.85% | 53.91% | 69.15% | 46.09% | N/A | N/A | N/A | N/A |
|  | ITDetector | 13.54% | 40.87% | 86.46% | 59.13% | N/A | N/A | N/A | N/A |
|  | Pindel | 5.00% | 15.65% | 95.00% | 84.35% | N/A | N/A | N/A | N/A |
|  | SoftSV | 16.24% | 16.52% | 83.76% | 83.48% | N/A | N/A | N/A | N/A |
| Tandem Duplications | MiStrVar | 100.00% | 86.67% | 0.00% | 13.33% | N/A | N/A | N/A | N/A |
|  | ITDetector | 3.17% | 80.00% | 96.83% | 20.00% | N/A | N/A | N/A | N/A |
|  | Pindel | 0.00% | 0.00% | 100.00% | 100.00% | N/A | N/A | N/A | N/A |
|  | SoftSV | 6.67% | 46.67% | 93.33% | 53.33% | N/A | N/A | N/A | N/A |

### 3.3 MicroSVs in the HCC1143 Breast Cancer Cell Line

Before running MiStrVar on the TCGA/CPTAC breast cancer samples, we applied it to the HCC1143 breast cancer cell line. We identified 116 microinversions and 197 microduplications (Supplementary Table 3) on this sample. Among these, 11 inversions and 12 duplications have both high read coverage and low mapping multiplicity. We focus only on these microSVs for the remainder of the discussion.

Details on the 11 inversion candidates can be found in Table 3. All 11 inversions appear in both normal and matching tumor samples indicating that they are germline events. 10 of them occur in intronic regions while one occurs in a 3’ UTR.

**Table 3:**
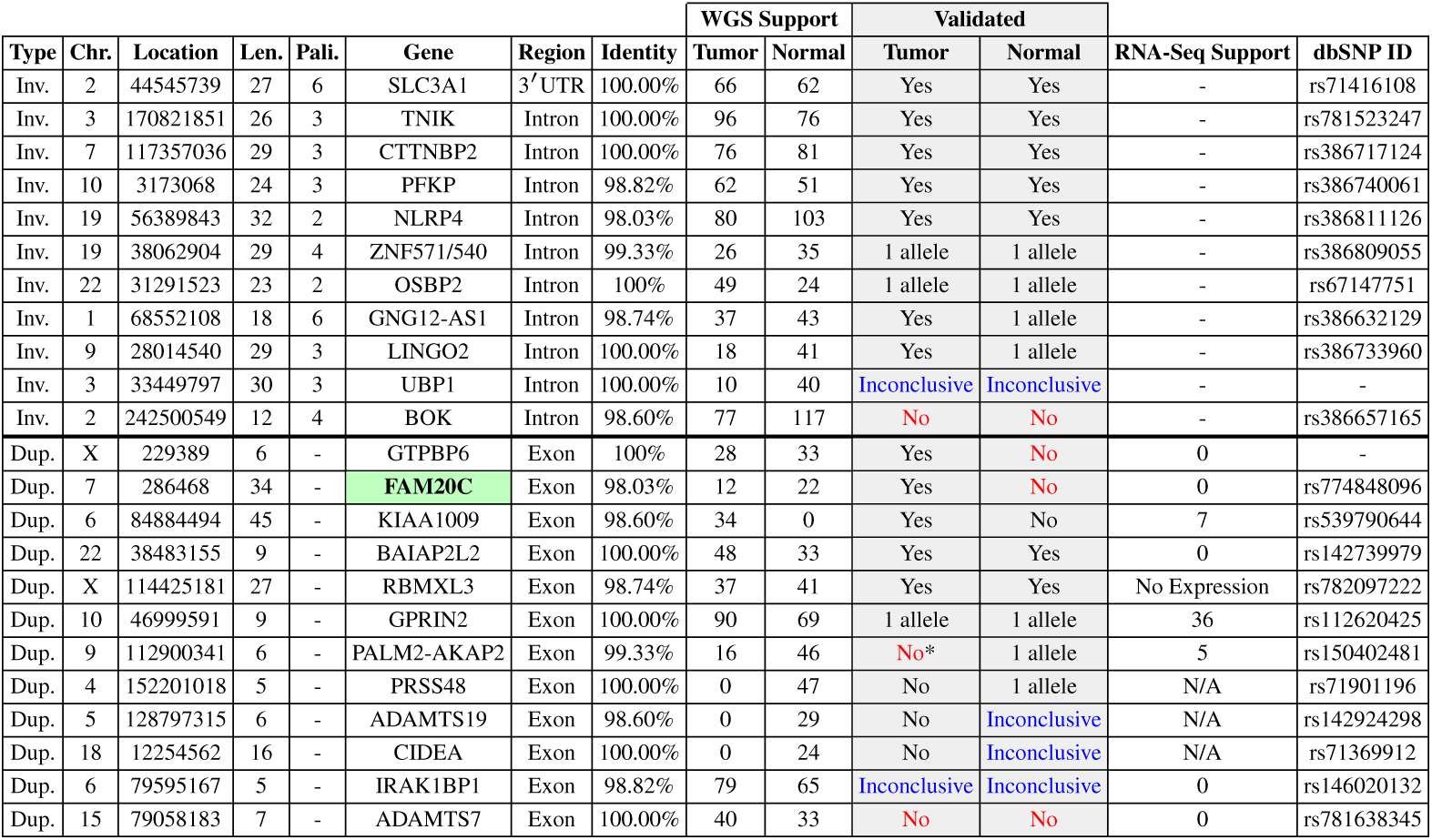
Sanger sequencing validation of top 11 microinversion and top 12 exonic microduplication (tandem or interspersed) candidates in the breast cancer cell line HCC1143. Entries marked “Yes” indicate a detected amplicon exactly matching the predicted microSV. “1 allele” indicates that two peaks were observed at each position in the chromatogram, only one matching the predicted microSV, and the other matching the reference, implying heterozygosity. For each detected inversion exactly matching an “multiple nucleotide polymorphism” and duplication exactly matching an “insertion” in dbSNP, we provide the dbSNP entry in the last column. As can be seen, all but two of these microSVs have been misclassified as a multiple nucleotide polymorphism or novel insertions in dbSNP. All microduplications are tandem, except for GTPBP6 which is interspersed. “RNA-seq support” denotes the number of reads support the structural variant. Since only tumor RNA-seq data was available, those SVs predicted in the normal sample are marked as “N/A”. The gene RBMXL3 is not expressed in this cell line therefore no supporting reads can be expected. Note that all of the microinversions we detected (with minimum support) were intronic and thus had no matching RNA-Seq reads. The duplication in PALM2-AKAP2 was likely missed by Sanger Sequencing in tumor (marked with an asterisk). The breast cancer-related gene FAM20C is marked in green.

We experimentally validated these inversions using Sanger sequencing. The primers were constructed by using the inverted sequence flanked by 200-300 bp from the reference genome. Five of the predicted inversions show a clear sequence match between the amplicon (from Sanger sequencing) and predicted inversion, validating these inversion candidates. A representative example is given for the inversion in SLC3A1 in Supplementary Table 7 and the complete set of chromatograms is included in the appendix. Four of the remaining inversions had amplicons with some nucleotides matching the reverse genomic strand and some matching the forward strand. This occurred in the amplicons from all four normal samples and two of the tumor samples. To resolve this discrepancy, the chromatogram corresponding to each amplicon was examined, first for the four normal samples, for which each of the inversion locations had either one or two peaks. In locations with two peaks, the bases always matched either the forward or reverse strand, exhibiting a classical case of heterozygous inversion that only occurs on one allele. For the final two inversion predictions, the amplicons for BOK and UBP1 corresponding to the tumor sample, only matched the forward genomic strand, which indicates no inversion at these locations. The amplicon corresponding to the normal sample of UBP1 contained many N bases in the sequence. Not enough information could be drawn from the chromatogram to conclusively say whether the amplicon supports an inversion.

We note here that all the high confidence microinversions, except for the one found in UBP1, have an associated multiple nucleotide polymorphism (MNP) entry in dbSNP. This includes the microinversion in BOK, which was not validated by Sanger sequencing.

In addition to MiStrVar we ran all the SV callers we tested on the HCC1143 cell line data. The parameters for all tools were identical to those used in the simulation. Out of these tools, only Pindel was able to identify any of the inversions. However, Pindel missed 2 of the 9 PCR validated inversion calls (in PFKP and OSBP2), out of 11 tested. The two calls made by MiStrVar that could not be validated were also called by Pindel, providing further evidence that MiStrVar improves Pindel with respect to both precision and recall.

The 12 duplication candidates are summarized in Table 3; all were exonic, i.e., fully or partially overlapping with exons. All of these duplications produced amplicons except for the one located in IRAK1BP1. Additionally, two amplicons from the normal sample (on genes ADAMTS19 and CIDEA) yielded a weak signal in the chromatogram so it was impossible to determine if they support the call or not; furthermore, the corresponding amplicon from the tumor sample showed no evidence of the call. Three of the nine remaining calls, in FAM20C, GTPBP6 and KIAA1009, show a clear match in the tumor sample but not in normal, indicating they are true somatic calls. Two calls, in BAIAP2L2 and RBMXL3, have a clear match in both tumor and normal samples, indicating they are germline calls. The next three showed two peaks at the insertion site and immediately downstream. One of the two peaks support the reference and the other the inserted sequence and the shifted reference, indicating that these calls are heterozygous. This was observed in both normal and tumor samples for GPRIN2 and only in normal for PALM2-AKAP2 and PRSS48. The final amplicon for ADAMTS7 showed only reference sequence at the insertion site, indicating that there is no duplication.

As per the microinversions, we ran all other computational tools mentioned earlier in order to determine if they are able to predict the validated microduplications. None of the tools were able to predict any of the microduplications. (Note that ITDetector was never able to complete execution after more than a month of processing.)

#### 3.3.1 MicroSVs in the Complete Set of TCGA-CPTAC Breast Cancer Samples

We applied MiStrVar and ProTIE to the complete set of matched tumor/normal samples from 22 TCGA breast cancer patients for which matching WGS, RNA-Seq and CPTAC Mass Spectrometry data were all available (see supplementary file for details). Minimal filtering was used on the calls since few calls uniquely matched proteomic signatures. Note that we only focus on exonic microinversion and microduplication calls (either fully or partially overlapping with exons) for further analysis. The number of calls for each sample can be found in Supplementary Table 6. Although only exonic calls were used for further analysis, the highest confidence calls within intronic and UTR regions, with respective support of > 40 and > 10 (identity = 100%) were also collected (see Supplementary Table 7). We also provide the highest confidence microduplications without proteomic support (support > 40, identity = 100%) as well as somatic microduplications (see Supplementary Table 8).

### 3.4 ProTIE Proteogenomics Analysis of CPTAC Breast Cancer Datasets

CPTAC has produced global proteome and phosphor-proteome data for 105 TCGA breast cancer samples using iTRAQ protein quantification method. Samples were selected from all four major breast cancer intrinsic subtypes (Luminal A, Luminal B, Basal-like/triple-negative, HER2-enriched) [57]. Each iTRAQ experiment included three TCGA samples and one common internal reference control sample. The internal reference is comprised of a mixture of 40 TCGA samples (out of the 105 breast cancer samples) with equal representation of the four breast cancer subtypes. Three of the TCGA samples were analyzed in duplicates for quality control purposes.

Our data analysis indicates that a two-dimensional reversed-phase liquid chromatographytandem mass spectrometric (2D-LC/MS/MS) sample comprises of about 0.87 million MS/MS spectra (per mixture). When we search them against Ensembl Human protein database, about 0.38 million MS/MS spectra in a mixture are matched to at least one peptide under 1% false discovery rate. These spectra lead to 59,387 proteins (42,840 known, 6,250 novel, 10,026 putative) with some peptides being covered by at least one spectra. The remaining 0.49 million spectra (≈ 56% of the whole set) do not match to any protein in the Ensembl database.

ProTIE obtains the intersection between these (0.49 million) unidentified spectra and the aforementioned set of fusions with missed cleaved polypeptides, to obtain 3,150,502 potential fusion peptides from 105 breast cancer patients^3^ (see Figure 1). ^4^ ProTIE uses a similar workflow to identify potential microSV peptides; for this case 635,125 potential microSV peptides were obtained from 22 patients.

Based on the database search strategy mentioned in supplementary subsection 2.3, in each mixture, our first level analysis resulted in approximately 5,342 spectra (1% FDR) matching to fusion peptide sequences, and about 620 spectra matching to microSV peptide sequences. If a matched peptide is identical to a substring of any known protein in Ensembl database, the corresponding spectra is discarded so as to ensure that the peptide is novel. The remaining results thus consist of all mass spectra in a single mixture supporting novel peptides originating from high confidence sequence aberrations. For a specific mixture, we can extract all the genes and the corresponding patient(s) generating these translated aberrations based on deFuse and MiStrVar calls.

It has been argued in the literature that stringent class-specific peptide-level FDR estimates may be necessary for reporting novel peptides in proteogenomics studies [2]. In order to address this issue, for any search result provided from MS-GF+, we first cluster all peptide-spectra matches into known or novel categories based on their peptide sequences: a PSM is assigned to the known class if the peptide is a known peptide or the decoy sequence of a known peptide; otherwise it will be assigned to the novel class. We then recalibrate FDR for records in the novel class using original E-value from MS-GF+: a peptide *p* is assigned the best spectral E-value *E*(*p*) it can get from any records in the novel class. Given a PSM *M* with E-value *s*, we collect all PSMs in the novel class whose E-value ≤ *s*, and calculate the ratio of records containing decoy sequences as the new peptide-level FDR for *M*. In tables 4, 5, and 6, a checkmark in the last column (labeled Str FDR) indicates that the corresponding PSMs pass this more stringent class-specific peptide-level FDR under 1%.

**Table 4:**
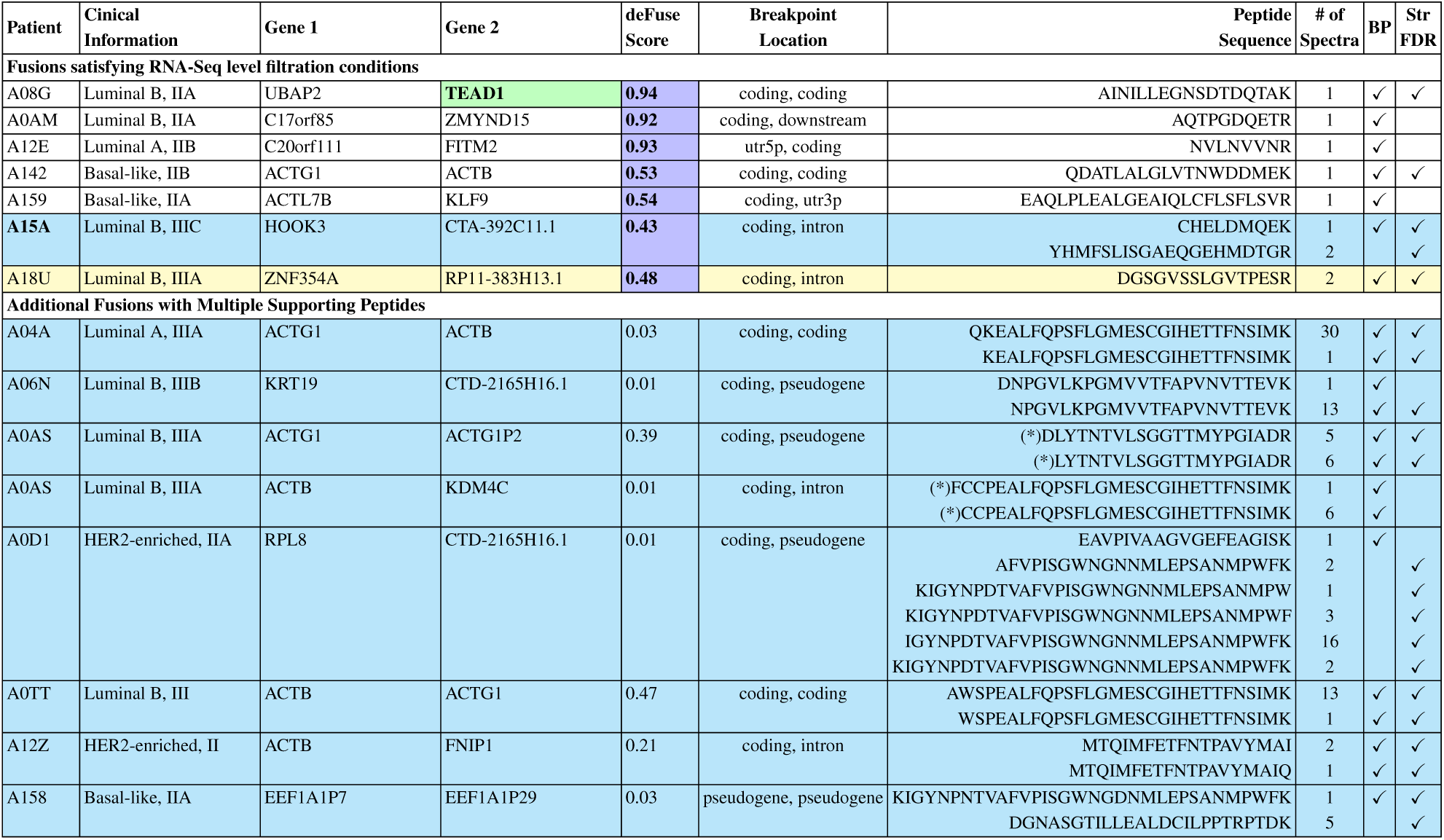
The list of selected (interesting) fusion events with translated peptides. A check mark in the column BP (BreakPoint) indicates that the peptide crosses the fusion breakpoint, and a check mark in the last column indicates that the peptide satisfies our more stringent FDR criterion. (a) *High confidence fusions*: fusions with high “deFuse Score” are colored purple (these satisfy stringent RNA-Seq level filtration conditions). (b) *Fusions with multiple supporting peptides:* fusion events associated with multiple novel peptides with proteomic support are colored cyan. (c) Among all fusions, one involves *a cancer gene*, TEAD1, and is colored green. (d) Only one fusion peptide is *supported by multiple spectra*: it is associated with the fusion detected in patient A18U, and is colored yellow. Note that peptides with star sign (*) are Single Amino Acid Variants (SAAVs) according to validated peptides in Ensembl GRCh38 protein database.

**Table 5:**
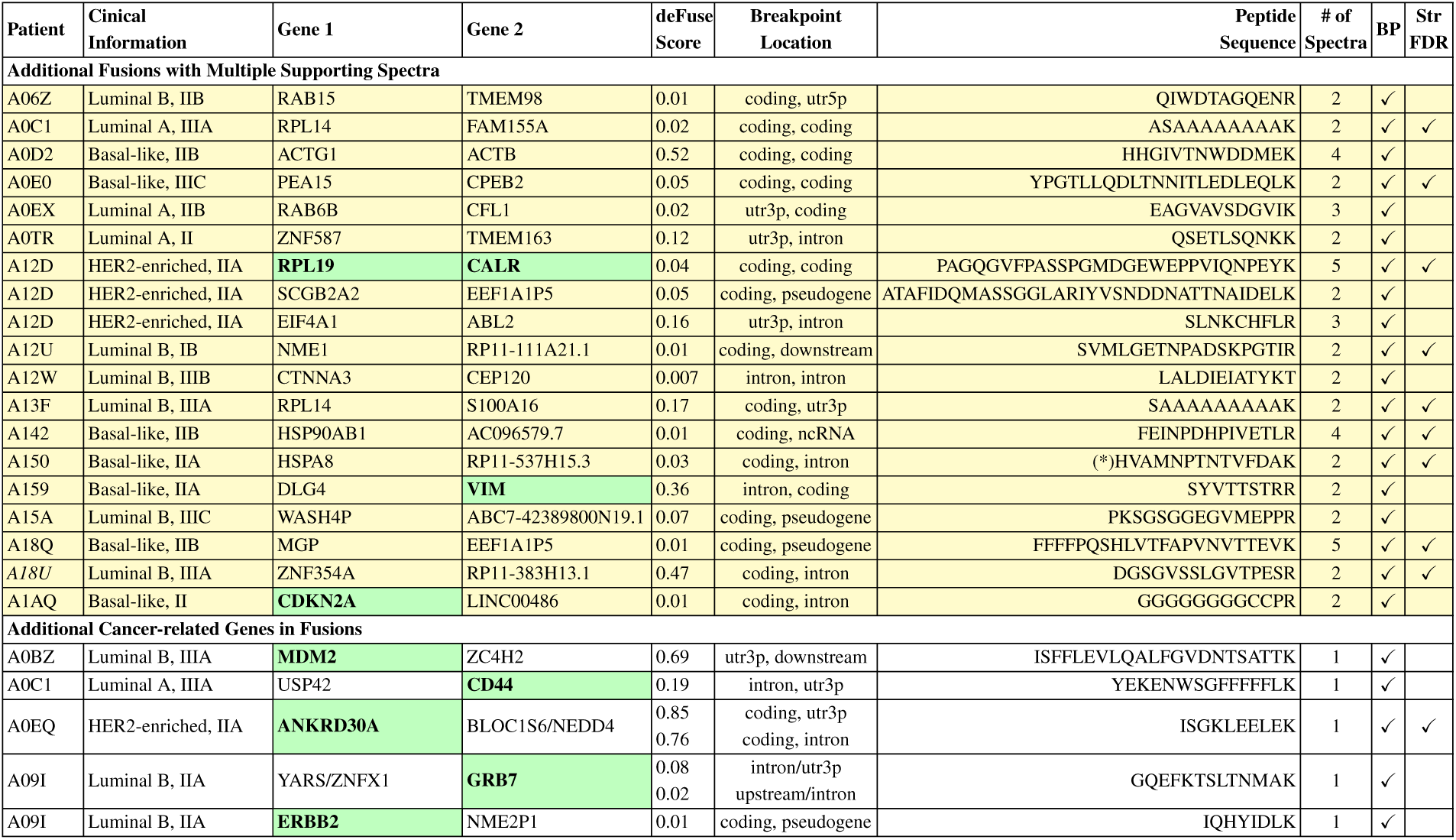
Additional list of selected (interesting) fusion events with translated peptides. A check mark in the BP (BasePair) column indicates that the peptide crosses the fusion breakpoint, and a check mark in the last column indicates that the peptide satisfies our more stringent FDR criterion. (a) *Fusions with multiple supporting spectra:* in addition to fusions in Table 4, other fusions have multiple supporting spectra - although all such spectra are associated with the same breakpoint-crossing peptide. These fusions are colored yellow. (b) *Fusions involving cancer genes:* fusions involving cancer-specific genes are colored green. Note that the peptide with a star sign (*) is a Single Amino Acid Variant (SAAV) according to validated peptides in Ensembl GRCh38 protein database.

**Table 6:**
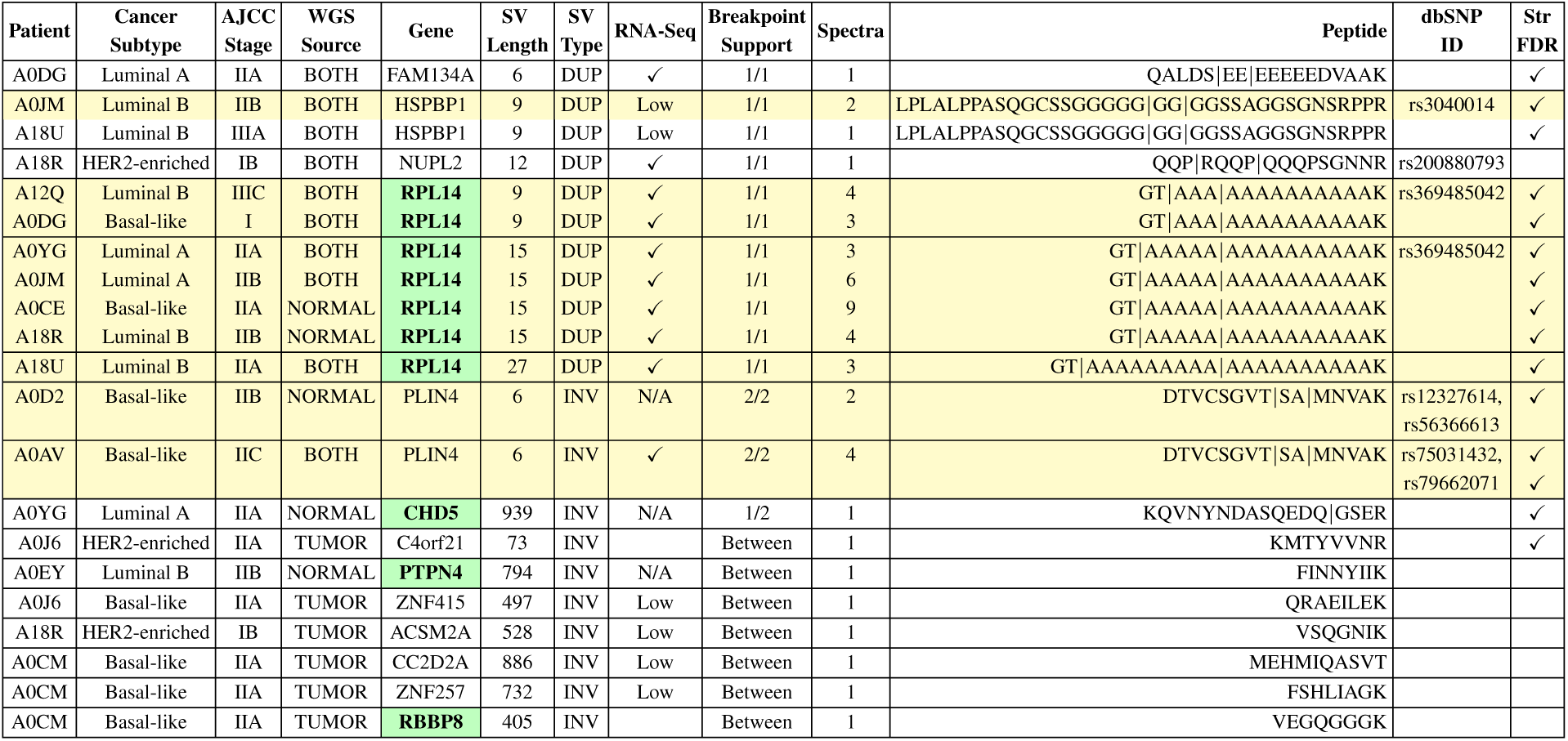
The list of genes containing microSVs with high confidence mass spectra support based on joint analysis of all 22 TCGA breast cancer patients with both tumor/normal WGS and tumor RNA-Seq data. For inversions, associated peptides always span one or two breakpoints (indicated as 1/2 or 2/2), or the inverted sequence between the breakpoints (indicated as “Between”). For duplications (which happen to be all tandem), the associated peptide always spans the single breakpoint and the entire inserted sequence (1/1). Breakpoints in the peptide sequences are marked with “|”. Calls marked as “Low” in the RNA-seq column are those from genes with low sequence coverage; similarly calls marked as “N/A” indicate the lack of RNA-seq data for this sample. Note that the microduplication in HSPBP1 is annotated as an insertion, and the microinversion in PLIN4 is annotated as two independent SNPs in dbSNP. Genes colored in green are known to be cancer related, and records colored in yellow have peptides with multiple supporting spectra.

#### 3.4.1 ProTIE Inferred Fusion Peptides

Given the proteomics search results for a specific mixture, a peptide will be further evaluated only if the corresponding fusion is also observed in at least one patient within the mixture. Among the remaining 5,579 spectra, 3,185 match to peptides coming from immunoglobulin heavy and light chain fusions. These peptides are not considered any further since highly repeated regions shared between those genes can lead to false positives in both fusion detection and proteomics search stages [58, 59]. Among the peptides remaining, we also discard those associated with a fusion for which no breakpoint crossing peptide is observed (This is due to the difficulty of determining whether such a peptide is a result of a fusion or because of a reading frame shift). At the end of these filtering steps ProTIE returns 807 spectra matching to 169 potential fusion peptides.

Among fusions related to these potential fusion peptides, we summarize special events with either high confidence RNA-Seq level evidence or proteomics support in Table 4. The first part of Table 4 shows events with better fusion quality based on reports of deFuse (deFuse score ≥ 0.1, cDNA percent identity < 0.1, EST and EST island percent identity < 0.3, no evidence detected for read through). Since 3 of these predicted fusions are between paralogs, specificially CRIP1 and CRIP2, IFITM2 and IFITM3, SRGAP2 and SRGAP2B, they are ignored. Among the remaining fusions, two stand out with respect to peptide-spectrum matching quality, respectively observed in patients A08G and A15A. The PSMs supporting these two fusions generated by pFind Studio [60, 61] are shown in supplementary materials.

We also provide a list of fusions with multiple translation peptides in the second part of Table 4. More specifically, four of these fusions have matching peptides located on both at the breakpoint and further downstream. Note that although we only detect a single peptide for some additional fusions, the peptide may be supported by multiple spectra as can be seen in Table 5.

#### 3.4.2 ProTIE Inferred MicroSV Peptides

As per ProTIE's translated fusion peptide inference approach, for each mixture, we only consider previously unknown peptides that can be unique byproducts of microSVs detected in at least one patient within the mixture. To ensure that these peptides support microSV (duplication or inversion) calls and not SNVs/SNPs, we only consider potential peptides from an interspersed duplication or inversion with a minimum of two amino acids on each side of at least one of the two breakpoints associated with that microSV; for tandem duplications we ensure that at least two amino acids are present in the peptide from both sides of the single breakpoint.

Proteomics search of these peptides on 22 patients resulted in 115 spectra potentially supporting microSVs. These spectra support a total of 75 peptides, due to the fact that some of the peptides are supported by more than one spectra. Of these 75 peptides, 7 support microduplications and 68 support microinversions. Incorporating the RNA-Seq results from section 4.3 in supplementary file, we obtain 4 microSV calls with support on all omics levels. The resulting peptides with the highest quality spectra support are summarized in Table 6. Here the number of spectra supporting these peptides is indicated in the “Spectra” column. Similarly, column “Breakpoint Support” indicates the number and type of the breakpoints supported by spectra for each peptide.

## 4 Discussion

### 4.1 Genomic MicroSVs Detected with MiStrVar

Our simulations show that MiStrVar effectively and accurately identifies all microSVs, specifically, insertions, tandem and interspersed duplications in WGS datasets. In particular, MiStrVar has high sensitivity, as well as high precision - especially for inversions, For duplications, even though its precision may not look as impressive, MiStrVar still outperforms all available alternatives. In addition, the precision values for duplications are likely to have been underestimated, since many of calls labelled as “false positives” could, in fact, be true germline differences between the Venter genome and the reference genome. On a very high coverage dataset (120x) from the Venter genome, with no simulated microSVs, duplications detected by MiStrVar have a large overlap with those it detects in the simulation dataset. Elimination of these calls from the simulation dataset increases MiStrVar's precision to 71% without any additional filtering.

MiStrVar is also very accurate in identifying the exact breakpoint loci of the microSVs. This is particularly important for our proteogenomics analysis where we only consider exact peptide matches. If a breakpoint were off even by only one nucleotide there is a high likelihood the predicted peptide would not match. With the exception of Pindel for inversions, which correctly identified 10% fewer exact breakpoints, no tool was even close to correctly identifying as many single-nucleotide resolution microSV breakpoints as MiStrVar. For inversions, the calls where MiStrVar can not identify the exact breakpoints are often due to the presence of palindromic sequences, resulting in co-optimal breakpoint predictions. More importantly, these cases yield identical peptides and therefore do not affect further analysis results. For duplications, the errors are usually observed in cases where the insertion is into a low complexity region. Again, in many of these cases the resulting peptides would be identical (e.g. consider a duplication that occurs in a polynucleotide tract). Furthermore, even in the worst case, MiStrVar predictions are within 30bp from the real breakpoints, still much better than the available alternatives. It should also be noted here that unlike other tools, MiStrVar provides not only the duplication breakpoint coordinates but also the precise coordinates of the “source” sequence (i.e. the region of the genome that is duplicated). Through this feature it becomes easier for the user to interpret interspersed as well as tandem duplications.

### 4.2 Translated Aberrations Detected with ProTIE

The use of a proteogenomic approach, as described in this study, enables two novel capabilities that are highly relevant to cancer biology and precision medicine. 1) The ability to hone in on potential clinically actionable mutations that are expressed at the protein level. The vast majority of clinical cancer testing focuses only on DNA-level mutations. A gene mutation-drug association is predicated on the assumption that a mutation will translate to the protein level, however, this is often not the case, as genes that contain a mutation may not be expressed in RNA. Moving further to the transcriptome the same paradigm exists, i.e., RNA expression does not always directly translate to protein expression, secondary to a variety of translational control mechanisms. Thus, having protein level evidence to confirm genomic aberrations provides assurance of the functional presence of a mutation. This has wide ranging implications for clinical cancer genomic testing, as well as the development of companion diagnostics for cancer targeted therapies. 2) The ability to observe the presence of protein spectra from fusion transcripts that are predicted to be out-of-frame. The vast majority of fusion annotation pipelines filter out fusions that are not in-frame secondary to a widely-held reasoning that these protein products would be misfolded and degraded or subject to non-sense mediated decay. Surprisingly, in this study, high quality spectra were observed from out-of-frame fusion spectra. While additional studies will need to be performed, these data suggest these out-of-frame fusion products are stable enough and at a relative abundance to be detected by Mass Spectrometry. Whether these products are stable by chance or confer a gain-of-function capability is yet to be seen, but these data at minimum suggest that out-of-frame fusions should not be eliminated from consideration (as is commonly done), when searching for oncogenic candidates.

#### 4.2.1 Translated Gene Fusions

To better understand the properties of genes with translation evidence for fusions, we analyzed these genes through Ingenuity Pathway Analysis (https://www.ingenuity.com). Note that we used all fusion genes detected by deFuse as the background genes in the analysis. The top 3 categories for gene function enrichment are: Cancer (137 genes), Organismal Injury and Abnormalities (150 genes), and Respiratory Disease (39 genes). All 3 sets of genes come with adjusted p-value around 0.0035 (via Benjamini-Hochberg procedure). Given that fusions are a somatic cancer-specific event, enrichment of cancer related genes provides a validation of our approach.

Many of the fused genes with detected novel peptides (each typically observed in a single patient) are associated with breast cancer. A selection of these fusions are listed in Table 4 and 5 where cancer-related genes are highlighed. Among them, a fusion of the Ubiquitin Associated Protein (UBAP2) and the transcriptional enhancing factor (TEAD1) is found in the patient A08G and meets our stringent FDR criterion. This fusion retains the DNA binding domain of TEAD1. Interestingly, high TEAD1 expression is associated with poor survival and this fusion may cause hyper-activation of TEAD1 in this patient [62, 63]. Note that the same fusion has also been detected with high confidence in TCGA Fusion gene Data Portal [64].

The remaining fusions associated with highlighted genes in Table 5 appear to be novel as they do not appear in the TCGA fusion database. Some of these fusions involve tumor suppressor genes. For example, even though the fusion detected in patient A0BZ does not meet our more stringent FDR criterion, it is interesting that it involves MDM2, a key regulator of the TP53 tumor suppressor pathway [65]. (TP53 is mutated in a large proportion of triple-negative breast cancers.) Another fusion that does not meet our more stringent FDR criteria but still is noteworthy is in patient A1AQ and involves CDKN2A gene, a tumor suppressor that inhibits the cell cycle and is deleted in many cancer samples [66]. The fact that it is fused to a long noncoding RNA, may be a novel mechanism to inactivate CDKN2A, as an alternative to deletion.

In addition to fused tumor suppressors, we also detected peptide evidence for fused oncogenes. The discovered fused oncogenes are: ANKRD30A, also known as NY-BR-1, a breast differentiation antigen observed in many breast cancer cells [67]; GRB7, a breast cancer driver gene which participates in Development ERBB-family signaling pathway [68, 69]; ERBB2, a well known breast cancer oncogene and biomarker [70] as well as the coexpressed gene Ribosomal protein L19 (RPL19); CALR, a gene highly expressed in approximately 5% of breast cancer cells and associated with metastasis [71]; and finally VIM, a protein involved in the epithelial to mesenchymal transition which drives metastasis [72]. The fusions involving ANKRD30A, RPL19 and CALR meet our stringent FDR criteria, while the others do not. In a number of cases, we can not pinpoint its fusion partners based on RNA-Seq data alone. The proteogenomics results help to increase our confidence of these fusions, and reduce the number of fusion partner candidates in the corresponding patients. The ERBB2 fusion is particularly interesting since ERBB2 is amplified in 15% of breast cancers and targeted with a variety of FDA approved drugs, making it a possible target for clinical analysis.

In the final list of 295 candidate fusions, 107 of the involved genes are also reported to be involved in a fusion according to TCGA Fusion gene Data Portal^5^. 58 of these genes have records in breast cancer (BRCA), and among them 19 genes are reported in the breast cancer database alone.

Among the ten cancer-related fusion genes in Table 4 and 5, nine are also found in TCGA Fusion gene Data Portal, with the exception of the ANKRD30A fusion. Seven of them (excluding VIM and CALR), are involved in fusions specifically in breast cancer patients. As mentioned earlier, UBAP2 is fused with TEAD1 in patient A08G, which matches the Fusion gene Data Portal entry exactly. The remaining six of these genes have different fusion partners in different patients.

#### 4.2.2 Translated MicroSVs

Most of the microinversions with proteomics support are in the 400bp to 1kb length range. Microinversions shorter than 100bp are much less common in exonic regions. However in intronic and UTR regions, microinversions with the best genomic support (in terms of both read coverage and sequence similarity - after the inversion is accounted) are predominately of length less than 100bp; See Supplementary Table 7, for a summary of intronic and UTR microinversions. We also observed that shorter microinversions tend to be germline events while longer events tend to be somatic.

All of the microduplication calls with proteomic support (all of these -with the exception of the one in NUPL2-satisfy our more stringent FDR criterion) were predicted to be germline events. Indeed nearly all of these events have corresponding dbSNP entries. The call in FAM134A appears to be a novel germline event. The longest duplication in RPL14 also appears to be novel (rs369485042 includes variants with up to 5 alanines). Deletions, translocations and allele loss at the genomic loci containing this gene has been observed in variety of cancers [73], including breast cancer [74]. This may be the case within patients AOCE, A18R (deletion) and A0JM (LOH). The unusually long case in patient A18U may lead to protein instability, causing the same phenotype as a deletion. Polyalanine tract lengths have been shown to be associated with cancer risk in other genes, such as androgen receptor in prostate cancer [75].

Since we observed relatively few translated microduplications, it is unlikely that this type of microSV plays a major role in breast cancer through translation to aberrant proteins. However we predicted many high confidence microduplications in exonic regions, some with RNA-Seq support, in addition to many in UTRs and introns (Supplementary Table 8). It is possible that such exonic duplications lead to truncated or rapidly degraded proteins and the duplications in UTRs and intronic regions may affect gene expression and splicing.

From our list of high confidence microSV calls (Table 6), four were found in genes known to be related to cancer (CHD5, RPL14, PTPN4 and RBBP8) and one in drug metabolism (CYP4F11). Among them, CHD5 is a particularly well studied tumor suppressor in neuroblastomas. It is also a known tumor suppressor in breast cancer [76], as well as colon, lung, ovary and prostate cancers [77]. The protein it codes, Chromodomain Helicase DNA binding protein 5, has functions in chromatin remodeling and gene transcription. CHD5 is frequently deleted in breast cancer and in one case a mutation resulted in a truncated, non-functional protein [76]. The microinversion we detected produces a stop codon shortly after the breakpoint which may also lead to the production of a truncated protein. Note that this microinversion satisfies our more stringent FDR criterion. Another interesting example, RBBP8 is a tumor suppressor specifically related to breast cancer. We have observed through inspecting geneMania [78] that RBBP8 is associated with the recombinational repair pathway (*p* < 1.27 × 10^−9^) (Supplementary Figure 11). RBBP8 is also known to modulate the important tumor suppressor BRCA1 [79] and act as a tumor suppressor itself through binding with the MRE11-RAD50-NBS1 (MRN) complex [80] or replication protein A (RPA) [81].^6^

Our analysis resulted in 4 microSV calls with support on all omics levels. This includes 3 microduplications (within genes FAM134A, NUPL2 and RPL14) and 1 microinversions (within PLIN4). The microduplications in FAM134A and RPL14 (that with 27bp) appear to be novel events. Additionally, there are several events with both genomic and proteomic support, which possibly lack RNA-Seq support due to low expression of the associated gene or the lack of RNA-Seq data for the sample.

## 5 Conclusion

Integration of genomic, transcriptomic, and proteomic data provides a comprehensive view of the patient's molecular profile. TCGA/CPTAC now offers matching genomic, transcriptomic and proteomic data across several cancer types, with a focus on the impact of Single Amino Acid Variants (SAAVs) and SNVs on protein abundances. In order to complement TCGA/CPTAC study and better establish the relationship between genomic, transcriptomic and proteomic aberrations and the cancer phenotype, we introduce MiStrVar, the first tool to capture multiple types of microSVs in WGS datasets. MiStrVar, and deFuse, a fusion detection tool we developed earlier, form key components of ProTIE, a computational framework we introduce here to automatically and jointly identify translated fusions and microSVs in matching omics datasets. Concurrently, ProTIE also incorporates RNA-Seq evidence to validate expressed microSVs. Based on both simulation and cell line data, we demonstrate that MiStrVar significantly outperforms available tools for SV detection. Our results on the TCGA/CPTAC breast cancer data sets also suggest the possibility of automatic calibration for some entries in dbSNP, which we believe are misannotated. It is interesting to note that the majority of the translated microSVs and fusions we observed in the breast cancer samples were private events; this prompts a larger and more detailed integrated study of all three omics data types through the use of ProTIE for a comprehensive molecular profiling of breast cancer subtypes.

## Acknowledgement

The WGS and RNA-Seq datasets were retrieved from the Cancer Genomics Hub (https://cghub.ucsc.edu/); the proteomics data was released by the National Cancer Institute Clinical Proteomic Tumor Analysis Consortium. Clinical information was obtained through the database of Genotypes and Phenotypes (dbGaP, http://www.ncbi.nlm.nih.gov/gap). Detailed description of the datasets used in this study can be found in https://wiki.nci.nih.gov/display/TCGA/TCGA+Data+Primer. We thank NIGMS/NIH Grant No R01 GM103725-04 (S.L.), and the NSERC Discovery Frontiers Grant on the Cancer Genome Collaboratory (C.S.) for funding this research.

## Footnotes

1 The primary goal of CPTAC is to characterize protein level expression differences for SNVs/SAAVs. Our focus here is complementary to the goals of CPTAC.

2 One interesting observation is that among the microSVs discovered, only 4 (specifically 1 microinversions and 3 tandem microduplications) have supporting evidence at all omics levels. This implies that the transcriptomic support for the remaining translated microSVs are too low to be detected, partially due to low abundance of RNA-Seq data made available by TCGA on the breast cancer samples we analyzed. This also suggests that with deeper coverage RNA-Seq data, ProTIE is likely to detect additional translated gene fusions.

3 Each breakpoint is associated with six reading frames and thus can result in (one of) six distinct proteins, and each such potential protein can lead to multiple potential peptides according to the number of K/R in the sequence.

4 Note that a reversed database was also appended here to control the false discovery rate.

5 Note that results in this database are based on 10431 calls from 2961 TCGA patients, which contains much broader scope than 105 breast cancer patients selected by CPTAC.

6 Binding of MRN and RPA occur through a domain at the N-terminus of the RBBP8 protein, which overlaps with the predicted microinversion. We hypothesize that the microinversion in this gene leads to the production of an aberrant peptide which is unable to bind to MRN or RPA, disrupting double stranded break repair and contributing to the cancer.

